# findere: fast and precise approximate membership query

**DOI:** 10.1101/2021.05.31.446182

**Authors:** Lucas Robidou, Pierre Peterlongo

## Abstract

**Motivation:** Approximate membership query (AMQ) structures as Cuckoo filters or Bloom filters are widely used for representing large sets of elements. Their lightweight space usage explains their success, mainly as they are the only way to scale hundreds of billions or trillions of elements. However, they suffer by nature from non-avoidable false-positive calls that bias downstream analyses of methods using these data structures.

**Results:** In this work we propose a simple strategy and its implementation for reducing the false-positive rate of any AMQ data structure indexing *k*-mers (words of length *k*). The method we propose, called `findere`, enables to speed-up the queries by a factor two and to decrease the false-positive rate by two order of magnitudes. This achievement is done one the fly at query time, without modifying the original indexing data-structure, without generating false-negative calls and with no memory overhead. With no drawback, this method, as simple as it is effective, reduces either the false-positive rate or the space required to represent a set given a user-defined false-positive rate.

**Availability:** https://github.com/lrobidou/findere

## 1 Introduction

Genomic studies generate a “data deluge” [13]. Public data banks providing sequencing data or assembled genome sequences are growing at an exponential rate [1]. Alongside, a fundamental need consists in comparing sequences at large scale, for instance for species identification [15], metagenome similarity estimation [3,11], or any generic need for estimating the presence of a sequence among available datasets [10].

Given the overwhelming amount of data to compare, these large scale sequence comparisons are made through alignment-free methods [16] mainly based on the number of words of fixed length, usually called *k*-mers for words of length *k*, shared between the two (set of) sequences to compare. So then, indexing *k*-mers for fast and low memory membership queries is a fundamental need. This last decade, an intense research activity was carried out in order to optimize such indexes (see [9] and [6] reviewing these efforts). Most of those indexes use approximate membership query (AMQ) structures such as Bloom filters [4] (called “BF” in this manuscript). With low space needed per represented element (usually less than 10 bits), these data structures are widely used although they suffer by nature from the existence of false-positive calls.

In genomics, BFs are the simplest and the most employed AMQ data structure used for representing the set of *k*-mers from a set of sequences. However, more sophisticated AMQ data structures enable under certain conditions to improve the BF features (cuckoo filters [7], SAT filters [14], quotient filters [2]). However, these improvements are marginal and/or at the expense of important limitations such as an important query time. For instance with SAT filters, despite they need ≈ 22% less space, their queries are roughly 14 times slower than BFs’ ones. Also, Cuckoo filters may show marginal smaller space cost per element for false-positive rates below 3% (space gain of 16% for instance for a false-positive rate of 0.1%, at the expense of longer query time).

In this work, we do not propose yet another AMQ data structure. Instead, we propose a downstream analyse of results returned by such data structures when used to query *k*-mers from a sequence. To the best of our knowledge, the only work in the same spirit is `kBF` [12]. `kBF` is designed to work on genomic sequences with the alphabet {*A, C, G, T*} and indexes *k*-mers using a BF. With `kBF` if a queried *k*-mer is positive in the BF, the presence in the BF of at least one of the four (the alphabet size) potential previous and incoming *k*-mers is also checked. If none of them is positive, then the original queried *k*-mer is considered a negative. This leads to a lower false-positive rate (up to 30x lower than a raw BF) at the expense of longer query time and higher memory usage. `kBF` presents interesting features that can be further improved as 1/ its query times are 1.3 to 1.6x longer than a classical BF, and 2/ its strategy is limited to the previous and the *k*-mer that comes just after a queried *k*-mer. Extending this approach to *n* neighbor increases the query time by (4^*n*^) fold. Finally, this approach applies only for small alphabets (eg. of size four) as the number of queries depends on the size of the alphabet. The work we propose aims to overcome these limitations.

In this paper, we propose a method for improving AMQ results when used for querying *k*-mers from a sequence. Our strategy is based on the observation that false-positive *k*-mers from AMQ are not likely to occur successively on a queried sequence. Hence, in a procedure that we call the “Query Time Filtration” (`QTF`), small stretches of positive calls surrounded by negative calls are considered as false-positives and are filtered out. This simple strategy leads to an unprecedented decrease of the AMQ false-positive rate. However, this leads to the introduction of false-negative calls that are a barrier for many downstream applications. Nevertheless, we show that the `QTF` strategy can be used for querying *K*-mers (with *K* > *k*), with no false-negative.

We implemented this approach in a tool called `findere`. Used on results from any original AMQ data structure, `findere` presents only advantages when querying *K*-mers: it does not necessitate any change to the original AMQ structure, it does not use any additional memory or disk, it has no false-negative calls, and it has a false-positive rate two orders of magnitude lower than original AMQ. Moreover, `findere` does not entail any additional query time penalty and even enables faster query of *K*-mers (in average > 2 times faster with recommended parameters) with no negative impact on result quality.

## 2 Method

### 2.1 Background

#### Preliminary definitions

A *k*-mer is a word of length *k* over an alphabet *Σ*. Given a sequence *∑*, |*S* | denotes the length of *S*.

In the current framework, we consider a dataset as composed of one sequence or a set of sequences. We consider that a *k*-mer occurs in a dataset if it occurs at least once in any of the sequences composing the set.

An AMQ data structure represents a set of elements *𝒟*. It can be queried with any element *d*. If *d* ∈ *𝒟*, then the AMQ answer is positive (there is no false-negative). If *d* ∉ *𝒟*, the AMQ answer may be negative or positive, in this last case it is a false-positive call. The false-positive rate, denoted by *FPR*_*AMQ*_, is defined by 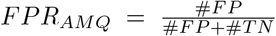 with #*FP* and #*TN* denoting respectively the number of false-positive calls and the number of true negative calls. *FPR*_*AMQ*_ depends on the used AMQ strategy and mainly on the amount of space used by this AMQ.

#### Sequence similarity estimated by the number of shared k-mers

Given two sequences, a *k*-mer occurring in both sequences is called “shared”. The number of shared *k*-mers between a queried sequence *q* and a dataset *B* provides an insight of the presence of *q* in *B*, or at least of a sequence similar to *q* in *B* [5].

In practice, for scaling Terabyte-sized sets *B*, a static AMQ indexes all (overlapping) *k*-mers from *B*. At query time, all (overlapping) *k*-mers from the query *q* are read on the fly and, for each of them, the *k*-mer is queried using the AMQ. Note that for each position *p* on *q* ∈ [0, |*q*| − *k* + 1] the *k*-mer starting at this position is queried. Thus, it overlaps by *k* − 1 characters with the previously queried *k*-mer if it exists (if *p* > 0).

### 2.2 Decreasing the AMQ false-positive rate with “Query Time Filtration”

In the context of computing the *k*-mer similarity between a bank *B* represented by its set of *k*-mers indexed in an AMQ and a query sequence *q*, we propose a surprisingly simple approach for drastically decreasing the AMQ false-positive calls.

#### Observation about the false-positive calls

Given the AMQ properties, one can assume that false-positive calls appear at random when querying elements from *∑*^∗^. In particular, when querying negative *k*-mers from a queried sequence *q*, the probability to query one false-positive is *FPR*_*AMQ*_, the probability to query two successive false-positives is *FPR*_*AMQ*_^2^, and so on. Overall, the probability to query *z* successive false-positives is *FPR*_*AMQ*_^*z*^. Given a classical *FPR*_*AMQ*_ value of 1% (usual expected false-positive rate with BFs for instance), with *z* = 3, the chances to call three consecutive false-positives *k*-mers is 0.0001%.

#### Intuition about the true positive calls

When approximating the sequence similarity between *q* and *B* using the *k*-mer similarity, the underlying idea is to choose *k* (usually chosen higher than 20 and lower than 40) such that *q* and sequences of *B* share large (≥ *k*) sub-sequences. An intuitive consequence of this choice stands in the fact that, when querying successive *k*-mers from *q*, it is unlikely that only a low number (*e*.*g*. less than 3) of successive *k*-mers are true positives.

#### Query time filtration

Motivated by the observation and the intuition presented in the two previous sections, we propose a method that we call the “Query Time Filtration” (`QTF` in short), designed for lowering the *FPR*_*AMQ*_ at the expense of the introduction of false-negative calls.

As illustrated Figure 1, the fundamental idea is to filter the AMQ answers depending on the context of the queried *k*-mers when those are queried successively. Given a parameter *z*, `QTF` answers “positive” only for *k*-mers having at least *z* consecutive neighbors indexed in the AMQ, and it answers “negative” else. Said differently, `QTF` considers as negative any *k*-mer that does not belong to a set of at least *z* + 1 successive *k*-mers positive for the AMQ.

**Fig. 1.**
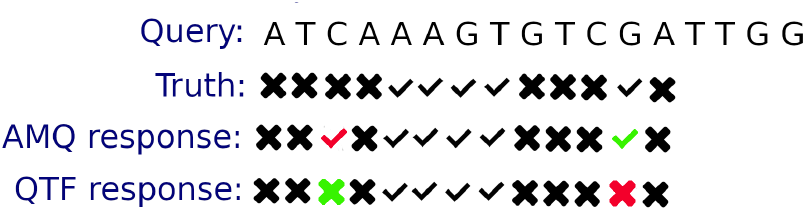
Example with *k* = 5 and *z* = 2, showing a query sequence (first line) and the various answers in the three next lines while querying 5-mers (indexed truth, AMQ answer and QTF answer). For each *k*-mer, a cross mark (resp. a check mark) indicates that this *k*-mer is absent from (resp. present in) the queried set. False answers are shown in red. This AMQ false-positive response (5-mer *CAAAG*) is filtered out by `QTF` as it generates a positive stretch of size one, lower than *z* = 2. Hence, one the last line showing the result after `QTF`, this false-positive does not occur. However, by applying this strategy, `QTF` may also remove the true positive stretch of size ≤ *z* (example shown with a red cross), leading to a false-negative QTF answer for the 5-mer *GATTG*.

In the following, a set of *x* consecutive *k*-mers positive for the AMQ is called a “positive stretch of length *x*”, or simply an “*x*-stretch”.

#### QTF algorithm

The `QTF` algorithm is straightforward. However, given its designed use cases (querying billions or trillions of *k*-mers) it has to be as much optimized as possible. We propose a simple yet efficient algorithm (see Algorithm 1) not using any extra disk space or RAM. Compared to a usual usage of an AMQ querying consecutive *k*-mers, it only requires a single additional integer “*currentStretchLength*” that represents the number of consecutive positives *k*-mers being read on *q* and a single test, with no impact on query time complexity that is *O*(|*q*|).

##### Algorithm 1

`QTF`

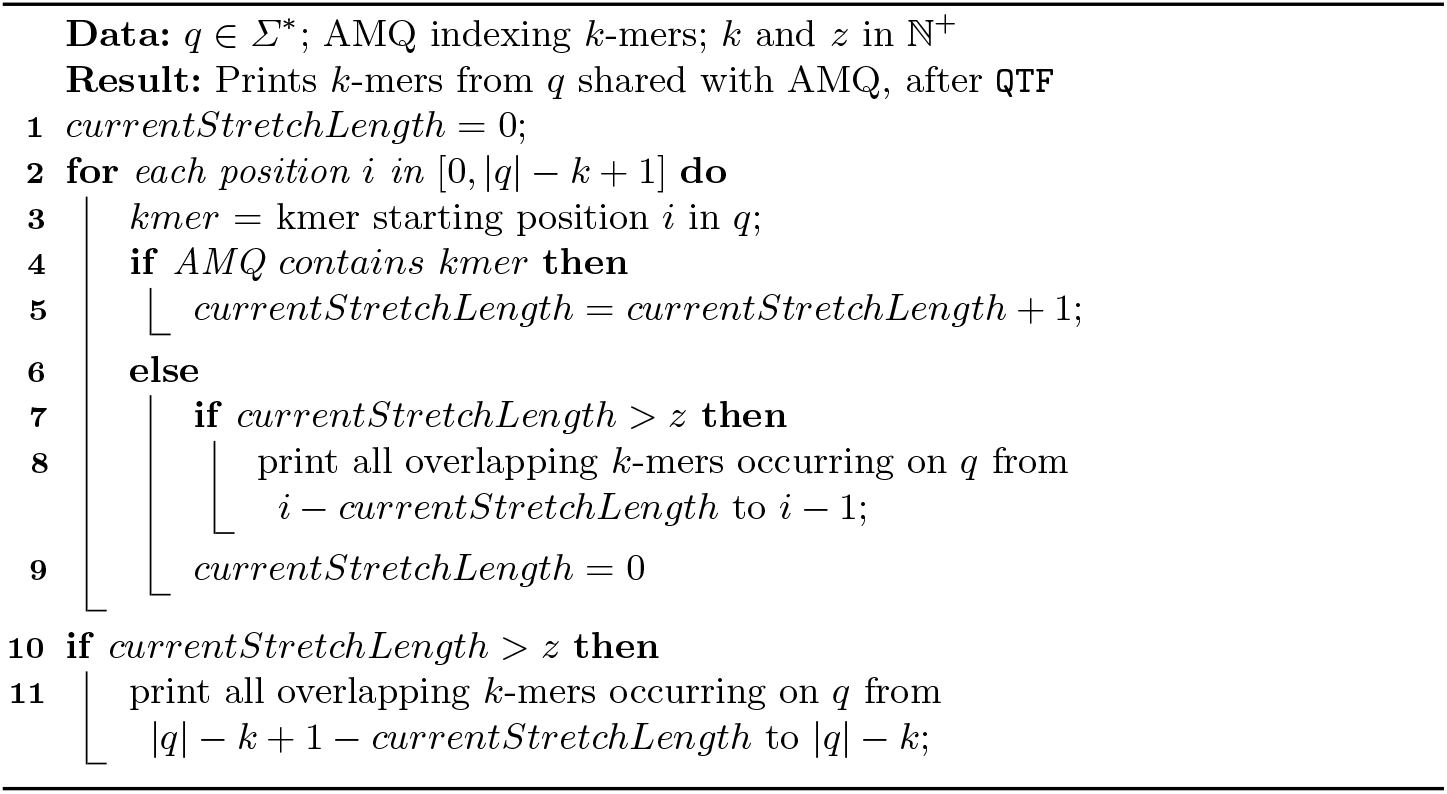

#### Considerations about QTF false-positive and false-negative rates

The `QTF` strategy drawback is the introduction of false-negative calls. A false-negative call occurs when a true-positive *k*-mer belongs to a *x*-stretch with *x* ≤ *z*. This situation happens when *q* and *B* share a (small) sub-sequence of length < *k* + *z*. This may happen in practice.

As shown by intermediate results (see Supplementary Materials, Section S1.1), the `QTF` strategy enables to reduce the original AMQ false-positive rate by several orders of magnitude at the expense of non-null false-negative rate. For instance, when applied to an AMQ composed of a BF indexing 31-mers with a false-positive rate of 5%, the `QTF` filter, using *z* = 3, enables to decrease the false-positive rate from 5% to ≈ 0.02% but increases the false-negative rate from 0% to ≈ 0.61%. With no impact on query time or memory, these results are highly satisfying for applications in which a small false-negative rate is acceptable.

By the way, in the following section we propose a second main contribution. We show how to take advantage of the `QTF` strategy to query *K*-mers (with *K* > *k*) with no false-negative, with low false-positive rate and with query time being faster than usual AMQ queries.

### 2.3 Querying *K*-mers with findere

As exposed previously, after `QTF`, a false-negative call arises only when a positive *k*-mer belongs to a positive stretch of length *≤ z*. Conversely, if a sequence of length *K* (with *K* > *k*) is shared between *q* and *B*, then it generates *K* − *k* + 1 successive true positive *k*-mers, thus defining a (*K* − *k* + 1)-stretch.

Conceptually, we take advantage of this remark by proposing an algorithm that we call `findere` in which we query *K*-mers based on indexed *k*-mers filtered by the `QTF` algorithm, using *z* = *K* − *k*, hence looking for stretches of length ≥ *z* + 1.

More precisely, given two integer values *K* and *k*, with *K* > *k* > 0, a bank dataset *B* and a query sequence *q*, the `findere` strategy consists in indexing all the *k*-mers from *B* using an AMQ. In contrast to `QTF` that calls *k*-mers, `findere` calls *K*-mers based on their *k*-mer content. Given a position *i* ∈ [0, |*q*| − *K* + 1] on *q*, the *K*-mer starting at this position is considered as “present” by `findere` if the *k*-mer starting at position *i* and the *K* − *k* next successive *k*-mers are considered as present by `QTF`. This explains the “ `findere`” name that comes from Latin and means “*divide*”.

The `findere` algorithm is obtained from a straightforward modification of the `QTF` algorithm (Algorithm 1). It is sufficient to define *z* = *K* − *k* and to print *K*-mers instead of *k*-mers lines 8 and 11, taking care to avoid printing the last *K* − *k K*-mers of each stretch. The complete algorithm is presented in Algorithm 2, including an additional time optimization as presented in the following section.

#### findere time optimization

While walking a sequence *q* searching for (*z* + 1)-stretches, it is possible to skip some *k*-mer queries. Indeed, if two negative *k*-mers start positions *i* and *i* + *z* + 1 on *q*, it is impossible to have a (*z* + 1)-positive stretch starting from any position between *i* and *i* + *z* + 1. This is because at most *z* positive *k*-mers can start between *i* and *i* + *z* + 1 both excluded, hence no (*z* + 1)-stretch can contain any position in [*i, i* + *z* + 1].

Thus, when a negative *k*-mer is found position *i* on *q*, we check directly whether the *k*-mer starting at position *i* + *z* + 1 is positive or negative for the AMQ.

– If it is negative, we know that no *z*-stretch can exists including any position in [*i, i* + *z* + 1]. There is in this case no need to query *k*-mers starting at

#### Algorithm 2

`findere`

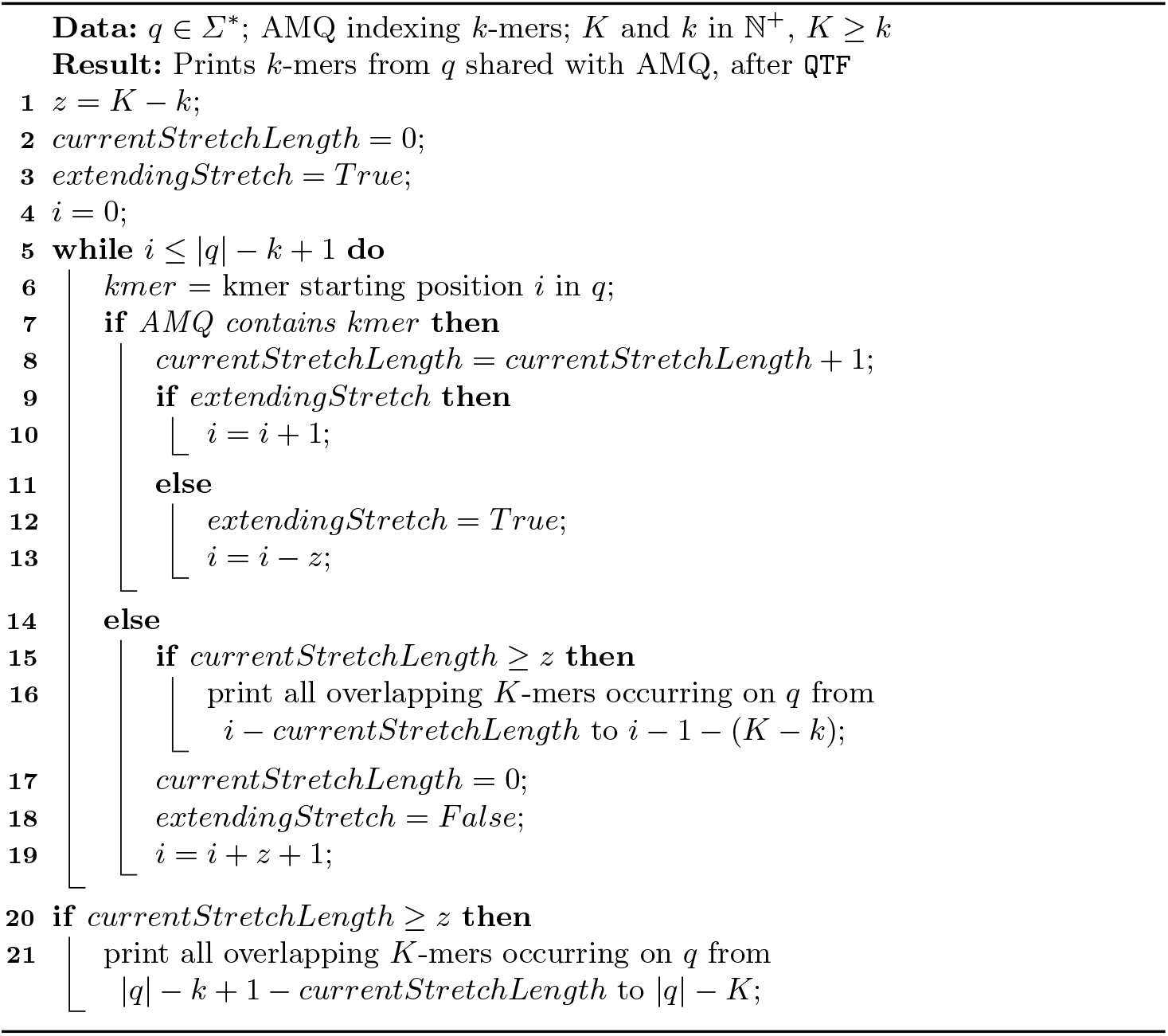 positions in [*i, i* + *z*], and we can repeat the process position *i* + 2(*z* + 1), and so on.
– If it is positive, *k*-mers starting from position *i*+1 have to be queried following the same process.

Note that even if we report this optimisation specifically for the `findere` algorithm, it also applies for the `QTF` strategy.

#### findere algorithm

Algorithm 2 proposes an overview of the `findere` algorithm. This includes the time optimization described in the previous section. This optimisation mainly takes effects line 19 where *z* positions are not queried unless the next checked position contains a positive *k*-mer, leading to rewind the query *z* positions back (line 13).

#### findere false-positives and “construction false-positives” (cFP)

`findere` reduces the false-positive rate of an AMQ by detecting and filtering out stretches of length ≤*z* (= *K* − *k*). The greater the length of the stretches, the fewer false-positives stretches pass through the filtration, as illustrated by results presented in Supp. Mat, Section S1.1.

However, detecting *K*-mers based on their *k*-mer content leads to the apparition of a novel kind of false-positives for `findere`. It may appear that for a negative *K*-mer, all its *k*-mers are true-positives. In this case, `findere` generates a false-positive call for this *K*-mer. We name those false-positive *K*-mers “construction false-positives” (cFP). The higher *z*, the higher the number of cFP, as when *k* gets too small, more *k*-mers occur by chance.

#### findere implementation

We propose an implementation of `findere`, available at https://github.com/lrobidou/findere. This implementation uses a Bloom filter as its inner AMQ. However, any other AMQ implementation can be used through a simple wrapper (provided with the `findere` implementation). The BF chosen for this implementation is a fork of the original https://github.com/mavam/libbf, which was modified to add the support of serialization. Although `findere` can index and query any alphabet, its implementation proposes a specialisation for genomic sequences: as such, one can index not only natural language, but also fasta and fastq files (gzipped or not) representing genomes or any sequencing read files. In this genomic context, a function to index and query canonical *K*-mers is also available.

## 3 Results

We propose results on real biological data and on natural texts. The aim is to show the practical advantages offered by `findere`, both in terms of query precision, index size, and query time. Being developed to be used on top of any AMQ, we do not compare `findere` with any of those. Remind that `findere` may be used on filtering the results from any such data structure, including Cuckoo Filters for instance. To the best of our knowledge, the only tool comparable to `findere` is `kBF`. We compared `findere` and `kBF` on biological data only as `kBF` is not designed to work on a generic alphabet.

Executions were performed on the GenOuest platform on a node with 4×8-cores Xeon E5-2660 2,20 GHz with 200 Go of memory. A complete description of tool versions, data acquisition, command lines, and numerical results are provided in the Github repository https://github.com/lrobidou/findere/tree/master/paper_companion.

### 3.1 Experimental data

#### Metagenomic data

In order to measure the impacts of the `findere` algorithm on real genomic data, we used two HMP [8] fastq files, indexing reads1 from sample SRS014107 and querying reads1 from sample SRS016349, both downloaded from the NCBI Sequence Read Archive. Theses samples contain respectively 4.2 million reads of average size 92 characters and 2.3 million reads of average size 96 characters. We simply refer to this dataset as the “hmp” dataset.

#### Natural language data

In order to test the `findere` implementation on natural language, we used a dump of Wikipedia, from which we extracted two subsets overlapping with 10 Mio. They have each a size of 100 Mio, leading to about 10^8^ 31-mers each. We refer to this dataset as the “natural language dataset”.

### 3.2 Results on genomics data

In this Section we propose results using *K* = 31. As shown in Supp. mat. (Section S1.2), `findere` results are robust with this main parameter.

#### False-positive analyses

We obtained results with a classical BF, indexing SRS014107 and querying SRS016349 with *K*-mers of length *K* = 31. For all experiences the size of the used BF is ≈ 2.6 billion bits, leading to 5% FPR when indexing 31-mers. With the same BF size, `findere` was run varying the *z* value, with *K* = 31. As shown Figure 2, with *z* = 0, `findere` obtains as expected exactly the same results as those obtained with the original used AMQ. With low *z* values (*e*.*g*. lower than 5), the `findere` FPR quickly drops close to zero. For instance, with *z* = 3, the `findere` FPR is equal to 0.056%, which is two orders of magnitudes smaller than the original BF FPR. Also, with such low *z* values, *k*-mers are large enough to limit the `findere` “construction FP” (cFP) to negligible values. For instance, *z* = 3 leads to *k* = 28 and the cFP rate is 0.025%.

**Fig. 2.**
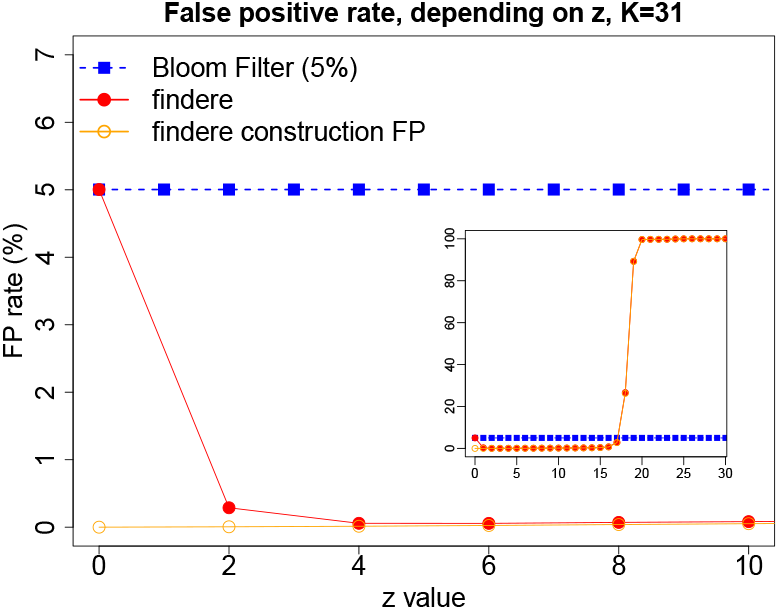
Comparative false-positive rate obtained on the hmp dataset. A BF indexes 31-mers, with an FP rate of ≈ 5% (blue filled squares, independent of the *z* value). With the same BF size, `findere` was run varying the *z* value, leading to FP rates as shown in red filled circles. Orange empty circles show the amount of `findere` “construction FP”, the rest of `findere` FP being due to stretches of length *≥ z* + 1 containing BF FP. The full figure zooms on recommended *z* values (in particular *z* ∈ [2, 5]). The small frame shows results including higher but discouraged *z* values.

When using large values of *z* (here > 15), *k*-mers get too small to be specific enough. Hence they have high chances to appear at random, leading to a dramatic increase of the cFP rate. This happens with *z* values leading to *k*-mers of size *<* 13. For instance, with *z* = 19, one has *k* = 12, which results for `findere` in an FPR of 89.27%, being almost only composed of cFP (curves overlap on the figure, cFP representing 99.95% of the FP).

Fortunately, fixing *z* is easily anticipated by choosing a value small enough so that the indexed *k*-mers in the AMQ have a low chance (*e*.*g*. lower than 0.01%) to appear by chance. The probability for a *k*-mer to appear by chance in a random text of length *n* can be roughly approximated as 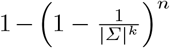, with |Σ| being the size of the alphabet. For instance on the indexed SRS014107 sample, where |Σ|= 4 and *n* = 386 millions, the probability to appear by chance for a *k*-mer is respectively equal to 0.99% for *k* = 13 (*z* = 18), to 0.30% for *k* = 15 (*z* = 16), to 0.02% for *k* = 17 (*z* = 14), and to 0.005% for *k* = 17 (*z* = 13). Hence choosing *z <* 14 is acceptable. By default `findere` uses *z* = 3.

#### Space gain

We recall first that `findere` memory usage has no overhead compared to the size of the used AMQ. For a given AMQ size, we can deduce the FPR for a BF and `findere`. Results are presented Figure 3 (with the default *z* = 3 value). This can also be used for deducing the amount of space needed for a given FPR. For instance, to obtain a usual value of 1% FPR, `findere` requires 0.05 Go of space while a BF requires 1.06 Go. The `findere` advantage gets even more important with a lower FPR: with 0.1% of FPR, `findere` requires 0.16 Go while a BF requires about 17 Go (dotted lines). This leads to a gain of space of two orders of magnitude, while not requiring any additional run time or RAM.

**Fig. 3.**
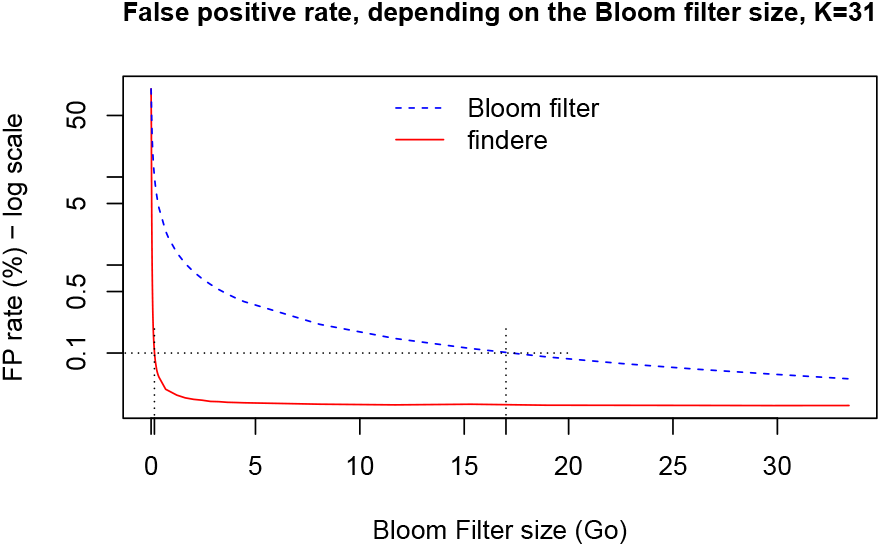
`findere` and BF FPR depending on the space used, on the hmp dataset. Dotted line segment corresponds to 0.1% false-positive rate.

#### Query time

Thanks to the optimization detailed section 2.3, the query time decreases when *z* increases. As shown Table 1, with discouraged values *z* = 0 or *z* = 1, the query time of `findere` is slightly higher than querying the original BF. This is due to additional conditional tests. With recommended *z* values (*z* = 2 to 5), compared to the query time of the BF, the `findere` query time is divided by a factor 2 to 3. With the default *z* = 3 value, query time is divided by 2.4, while query time still decreases when *z* increases: with *z* = 10, the query time is divided by ≈ 5.

**Table 1.** BF and `findere` query time in seconds on the hmp dataset, depending on the *z* value. BF result does not depend on *z* and is reported only for *z* = 0.

| $z$ | 0 | 1 | 2 | 3 | 4 | 5 | 10 |
| --- | --- | --- | --- | --- | --- | --- | --- |
| BF | 42.4 |  |  |  |  |  |  |
| **findere** | 42.9 | 43.7 | 24.3 | 17.5 | 14.1 | 12.0 | 8.6 |

#### Comparisons with kBF

To compare `findere` and `kBF`, we created a fork of `kBF`, available at http://github.com/lrobidou/kbf/. This fork enables to specify the amount of memory to be used by `kBF` and more importantly, it enables to index a set of reads and query another set (the original implementation does not allow that). `kBF` comes with two versions: 1-`kBF` and 2-`kBF`. 1-`kBF` uses less space at higher FPR than 2-`kBF`. The `kBF` strategy imposes to query up to nine times the BF when asking for the membership of a single *k*-mer. At the same time, `findere` is ≈ 2.4 times faster than a Bloom filter with the recommended value *z* = 3. Moreover, `kBF` dumps all queried *k*-mers in RAM, and 2-`kBF` uses an additional hash set. Hence, `kBF` is much slower than `findere`. For instance, on the hmp data, with a BF FP of 5%, 1-`kBF` (resp. 2-`kBF`) query needs ≈ 300s (resp. ≈ 1450s) while `findere` needs ≈ 17s.

Moreover, as it shows a higher FPR for a fixed amount of space, `kBF` uses more space than `findere` for an equivalent FPR. For instance with for ≈ 1% FPR, `findere` requires 0.05 Go of space while 1-`kBF` requires ≈ 0.40 Go. Comparisons with 2-`kBF` are not fair as it computes a hash set of every *k*-mer, leading to unreasonable space usage (*e*.*g*. 7.78 Go for an FPR of ≈ 1% when `findere` requires 0.05 Go). Full `kBF` results are proposed in Supp. Mat, Section S1.4.

### 3.3 Results on natural languages

We applied `findere` and BF on the natural language corpus. Full results are provided in Supp. Mat, Section S1.3.

#### Memory gain

As in Section 3.2, we computed the FPR in function of the space used. From those results, we can deduce that, also on natural languages, for an FPR of 0.1%, the `findere` space usage is two orders of magnitude less than BF. Indeed, `findere` needs 0.023 Go, while a BF requires 3.38 Go.

#### Query time

As described in section 2.3, the query time decreases when *z* increases. It holds when `findere` is used on the natural language dataset as well. With recommended *z* values (*z* = 2 to 5), compared to the query time of the Bloom filter, the `findere` query time is divided by a factor 1.6 to 3. With the default *z* = 3 value, query time is divided by 2.2 compared to the raw Bloom filter query.

### 3.4 Limit of the findere approach

The `findere` algorithm generates so-called “construction false-positives” that occur when a negative *K*-mer contains only true positive *k*-mers. With recommended parameters, those cFP are negligible as shown in Fig. 2. However, as cFP depends only on true-positive calls, its value does not depend on the *FPR*_*AMQ*_. Hence theoretically, when *FPR*_*AMQ*_ tends toward zero, the cFP rate (and thus `findere` FPR as well) becomes higher than *FPR*_*AMQ*_. This effect can be observed on the natural language results (Fig. S2, Supp. Mat.) with BF FPR below 0.02%. However, one should remind that, first, the difference is insignificant (0.008% difference FPR when using 26 Go space), and second, the practical usage of an AMQ is usually with FPR higher than 0.1% to prevent huge space requirements.

## 4 Conclusion

We propose a method filtering results of any approximate member query (AMQ) data structure when used for querying words of length *K* from a query. Despite its amazing simplicity, applied on metagenomics and natural text data, compared to the non-filtered results: `findere` **1/** makes queries two times faster, **2/** enables to decrease by two orders of magnitude the false-positive rate or enables to decrease the space allocated to each element by two orders of magnitudes, and **3/** has no drawback when used with recommended values.

We are expecting an important impact of the `findere` tool for which we propose an implementation. Indeed, AMQ data-structure are essential for indexing large datasets. In particular their usage is fundamental for indexing the genomic sequencing data for which `findere` offers a new scaling breakthrough.

## Acknowledgements

This work used HPC resources from the GenOuest bioinformatics core facility (https://www.genouest.org). The work was funded by ANR SeqDigger (ANR-19-CE45-0008).

## S1 Supplementary Materials

### S1.1 QTF false positive analyses

In this section, we show qualitative results obtained thanks to the `QTF` strategy that motivated the `findere` algorithm. Results shown Table S1 were obtained by applying the `QTF` approach, using *k* = 31, with various *z* values on an AMQ data-structure (a bloom filter) with 5% false positive rate (*FP*_*AMQ*_ = 5%). Indexed and queried data are those from the hmp project, as described Section 3.1.

**Table S1.** Qualitative results using the `QTF` strategy, applied on the hmp dataset with *FP*_*AMQ*_ = 5%.

| $z$ | #TP | #FP | #FN | #TN | FPR (%) | FNR (%) |
| --- | --- | --- | --- | --- | --- | --- |
| 0 | 4341814 | 5352422 | 0 | 101656172 | 5 | 0 |
| 1 | 9143324 | 540743 | 10169 | 101656172 | 0.529 | 0.111 |
| 2 | 9600802 | 58782 | 34652 | 101656172 | 0.058 | 0.360 |
| 3 | 9613606 | 21273 | 59357 | 101656172 | 0.021 | 0.614 |
| 4 | 9585992 | 17880 | 90364 | 101656172 | 0.018 | 0.934 |
| 5 | 9540667 | 16935 | 136634 | 101656172 | 0.017 | 1.412 |
| 6 | 9500603 | 15976 | 177657 | 101656172 | 0.016 | 1.836 |
| 7 | 9456841 | 15296 | 222099 | 101656172 | 0.015 | 2.295 |
| 8 | 9396421 | 14630 | 283185 | 101656172 | 0.014 | 2.926 |
| 9 | 9341993 | 13827 | 338416 | 101656172 | 0.013 | 3.496 |
| 10 | 9284567 | 13265 | 396404 | 101656172 | 0.013 | 4.095 |

These results highlight the dramatic decrease of the false positive rate of the original AMQ data structure. Indeed, the false-positive rate falls from 5% with *z* = 0 where `QTF` has no effect, to 0.53% when using *z* = 1 (removing all 1-stretches). With *z* = 2, the false positive rate is again reduced by an order of magnitude. However, the false-positive rate, being null with the original AMQ raises to ≈ 0.1% with *z* = 1, and reaches ≈ 0.36% with *z* = 2.

### S1.2 Results varying *K*

We varied the value of *K*, and we report the false positive rate as well as the query time. Tests were performed on the hmp dataset using the default *z* = 3 value. For each *K* value, we determined *size*_*K*_ being the size of the BF such that the FPR of the BF for *K* is 5%. Then, for each *K* value, we computed the `findere` FPR and query time, using the *size*_*K*_ BF size for indexing *k*-mers. Results, also comparing results that would be obtained with the non filtered BF, are presented Fig S1.

**Fig. S1.**
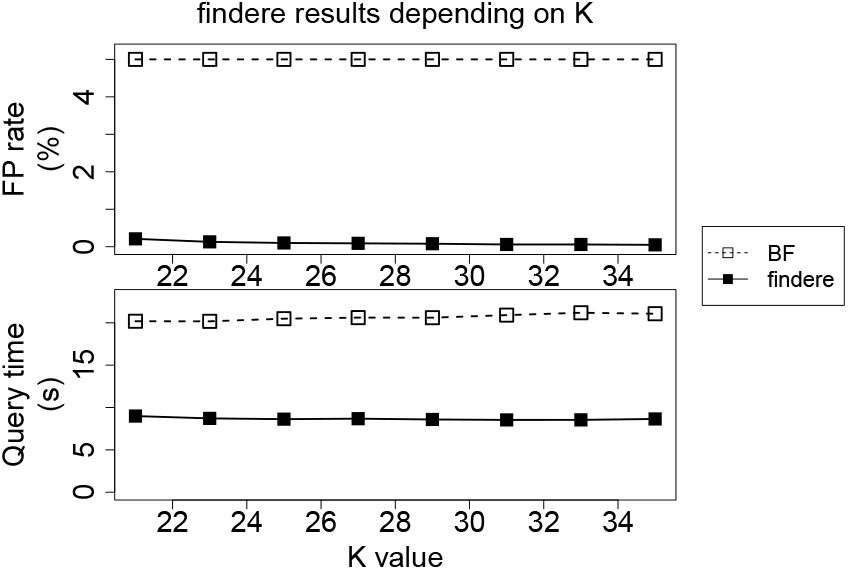
`findere` FPR and query time depending on *K*, on hmp dataset, using *z* = 3.

### S1.3 Results on natural languages

In this section, we show results obtained when applying `findere` on the natural language dataset. Figure S2 shows the FPR of `findere` and Bloom filters when indexing 31-mers from this dataset depending on the size of the index. Table S2 shows the time taken by a Bloom filter and `findere` to query this dataset with respect to the value of *z*.

**Table S2.** BF and `findere` query time in seconds on the natural language dataset, depending on the *z* value. BF result does not depend on *z* and is reported only for *z* = 0.

| $z$ | 0 | 1 | 2 | 3 | 4 | 5 | 10 |
| --- | --- | --- | --- | --- | --- | --- | --- |
| BF | 19.36 |  |  |  |  |  |  |
| **findere** | 19.87 | 19.99 | 11.76 | 8.77 | 7.32 | 6.56 | 5.56 |

### S1.4 Comparison with kBF on the hmp dataset

To date, in the `kBF` implementation, the *k*-mers are read from a file, stored in a vector, and indexed. A fraction of these *k*-mers is mutated before querying all the stored *k*-mers. Thus, the `kBF` implementation does not propose a way to index a set of reads and to query another set of reads. Furthermore, its implementation does not enable to serialize the index, and does not enable to set or get the size of the index.

**Fig. S2.**
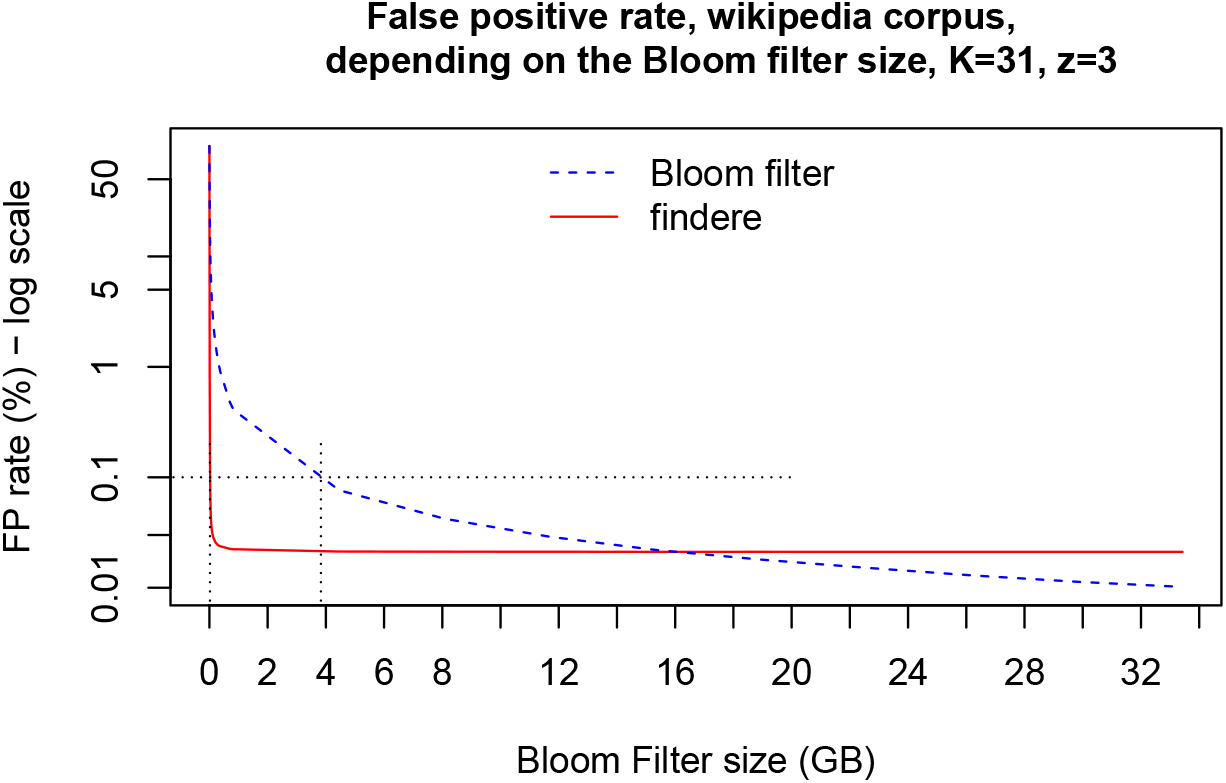
`findere` and BF FPR depending on the space used by their index, on the natural language dataset. Dotted line segment corresponds to 0.1 % false positive.

Finally, it is not possible to pass directly the amount of space to `kBF`. Instead, the user must provide an integer value (called “size factor”), which is the desired number of bits per element in the bloom filter used by `kBF`. Consequently, the space used by its bloom filter is always a multiple of the number of elements indexed. The size factor is decided at compilation time for 1-`kBF` and cannot be changed later on.

As a consequence, we proposed a fork of `kBF` that allows to index a set of reads and query another one, as well as to change the size factor for 1-`kBF`. Then, using the size factor, we were able to determine space and memory 1-`kBF` would end up using. However, 2-`kBF` needs to store an additional hash set of *k*-mers. Since we were not able to get the amount of space taken by that hash set, we do not report it here. Consequently, the “size required” column in Table S3 is to a great extent underestimated for 2-`kBF`. Indeed, the peak RAM usage is between 7 Go and more than 8 Go for all experiments proposed in this Table.

From Table S3, we can observe that `findere` outperforms Bloom filters, 1-`kBF` and 2-`kBF`. For a high number of bit per element, the FPR of `findere` and 2-`kBF` seem close, however, the memory is heavily underestimated in the case of 2-`kBF`, and as such, 2-`kBF` is likely to take much more space (*e*.*g*. more than 8 Go when using 26 bits per element in the BF).

In Table S4, we show that, for the indexation and the query steps, both 1-`kBF` and 2-`kBF` are slower than the Bloom filter used in the implementation of `kBF`.

**Table S3.** FPR of Bloom filter, 1-`kBF`, 2-`kBF` and `findere` with respect to the size of the filter, for *K*=31, on the hmp dataset. `findere` was used with default *z* = 3 value.

| number of bits<br>per element<sup>1</sup> | size<br>required (Go)<sup>1</sup> | FPR<br>BF (%)<sup>1</sup> | FPR<br>1-kBF (%) | FPR<br>2-kBF (%) | FPR<br>findere (%) |
| --- | --- | --- | --- | --- | --- |
| 3 | 0.05 | 28.34 | 26.38 | 15.40 | 0.95 |
| 5 | 0.09 | 18.12 | 14.49 | 5.52 | 0.27 |
| 7 | 0.12 | 13.32 | 9.10 | 2.54 | 0.15 |
| 9 | 0.15 | 10.52 | 6.22 | 1.37 | 0.11 |
| 15 | 0.26 | 6.45 | 2.69 | 0.36 | 0.07 |
| 18 | 0.31 | 5.41 | 1.96 | 0.22 | 0.06 |
| 21 | 0.38 | 4.65 | 1.49 | 0.14 | 0.05 |
| 24 | 0.41 | 4.08 | 1.17 | 0.10 | 0.05 |
| 26 | 0.44 | 3.77 | 1.01 | 0.08 | 0.05 |

<sup>1</sup>Not including the hash set for 2-kBF

**Table S4.** Time taken (in second) by `kBF` to index and query all of the 31-mers in the hmp dataset.

| size factor<sup>1</sup> | 3 | 5 | 7 | 9 | 15 | 18 | 21 | 24 | median |
| --- | --- | --- | --- | --- | --- | --- | --- | --- | --- |
| time index bf (s) | 223 | 235 | 239 | 242 | 248 | 249 | 250 | 250 | 245 |
| time query bf (s) | 178 | 194 | 202 | 207 | 214 | 215 | 217 | 218 | 211 |
| time index kbf (s) | 220 | 236 | 240 | 241 | 245 | 247 | 248 | 250 | 243 |
| time query kbf (s) | 329 | 342 | 340 | 332 | 316 | 309 | 304 | 300 | 323 |
| time index kbf2 (s) | 506 | 470 | 442 | 419 | 377 | 365 | 355 | 350 | 398 |
| time query kbf2 (s) | 1399 | 1420 | 1433 | 1440 | 1453 | 1453 | 1453 | 1475 | 1446 |

<sup>1</sup>The size factor only takes into account the bloom filter used by kBF, not the adjacent hashSet

## Notes

https://github.com/lrobidou/findere

### Competing Interest Statement

The authors have declared no competing interest.

### Summary of Updates

Correction of a sentence section 2.2, about the chances to call three consecutive false-positives k-mers.

